# The identification of two newly discovered fluorescent proteins in human glioblastoma

**DOI:** 10.1101/2022.03.15.484413

**Authors:** Xiao Lyu, Yutian Wei

**Author notes:** **Corresponding authors:** Yutian Wei, Department of Neurosurgery, The First Affiliated Hospital of Nanjing Medical University, Nanjing, 210029, China. These authors contributed equally to this work. **Consent for publication:** All co-authors have read and approved of its submission to this journal.

## Abstract

Glioblastoma (GBM) is a common malignancy featured by an extremely strong proliferation with enriched genetic information. Due to the rapid proliferation and the unstable genome-induced evolutionary potential, unique proteins may be expressed in GBM cells under the certain influence of the microenvironment. We therefore speculated that fluorescent proteins exist in GBM cells. During the immunofluorescence staining assay, we accidently discovered autofluorescence in primary GBM cells without fluorescent labeling, which were further validated as 2 newly discovered fluorescent proteins excited by 467 nm and 378 nm wavelength, respectively, namely human fluorescent protein Ⅰ and Ⅱ (HFP1, HFP2). Fluorescence colocalization and fluorescence resonance energy transfer (FRET) results showed the tight interaction of HFP1 and HFP2, and their synergistic effect. Our results for the first time identified 2 newly discovered fluorescent proteins in GBM cells, and clarified their chemical properties.

## Introduction

Existing evidences have shown that opsins and fluorescent proteins are the two types of photosensitive proteins in organisms. The former is found in ocular organs or bacteria, and the latter is only expressed in metazoans, especially in jellyfish, corals polyps (phylum Cnidaria), comb jellies (Ctenophora), crustaceans (Arthropoda) and lancelets (Chordata)^1^. For example, the chromophore of wild-type green fluorescent protein (wtGFP) is formed by the cyclization and dehydrogenation of the serine 65 (Ser65), tyrosine 66 (Try66), and glycine 67 (Gly67) residues in the α-helix. The conjugated π-electron on the wtGFP chromophore contributes to absorb the excitation energy, and immediately release it at a longer wavelength to form the fluorescence signal.Most of genomes in animals lack GFP sequences, suggesting that GFPs are originated in the early stage of evolution and deficient in many species^1^. So far, biological functions of GFP-like proteins have not been comprehensively analyzed, and relevant experimental data on them in many species remain largely unclear^1^.

Our research group has made great efforts on the research of GBM, and we were capable of culturing primary GBM cells. We directly isolated primary GBM cells from cancer specimens without cell passage before experiments, thus preserving cell primitiveness as much as possible. The evolutionary mechanism of GBM is similar to that of asexual reproduction species following replication, genetic variation, genetic drift, selection and environmental adaptation^2^. Hence, we speculated that primary GBM cells are a unicellular organism containing the human genome, in which the photon motion is involved as an environmental factor. GBM cells are responsive to abundantly express fluorescent proteins that accept photons and positively stimulate the disease progression, expression levels of which may vary a lot due to the cancer heterogeneity and individual differences.

In the present study, we first isolated primary GBM cells from clinical specimens of 6 GBM patients, and identified newly discovered HFP1 and HPF2. Their expression levels in different cell lines were detected.

## Methods

### Ethics approval and consent to participate

The study was approved by the Ethics Committee of Nanjing Medical University (code: 2019-SR-479) and written informed consent was obtained from all patients. The diagnosis of glioblastoma was confirmed by pathologists. Detailed patient information is presented in Table S1.

### GBM samples and primary GBM cells

Six GBM specimens were surgically collected in the Jiangsu Province Hospital, Nanjing Medical University (Nanjing, China). Briefly, 1 cm3 GBM tissues were washed in 300 µL of 1×PBS, which were cut into small pieces (1 mm3) and digested in a mixture containing 1 mL of 5 mg/mL neutral protease, 1 mL of 2 mg/mL collagenase, 3 mL of pre-warmed DMEM/F12 (1:1) and 30 µL of 1000 units/mL DNase. The precipitant was resuspended in DMEM/F12 (1:1) medium and cultured in an incubator at 37ºC containing 5% CO2 in the dark. Fresh medium was replaced with an interval of 4 days.

### Cell culture

GBM cell lines U251, U87 and U87-GFP, and human gastric mucosa epithelial cell line GES-1 (ATCC) were cultivated in DMEM (HyClone) containing 1% penicillin and streptomycin, and 10% FBS in an incubator at 37ºC containing 5% CO2.

### Confocal immunofluorescence microscopy

Cells were seeded in laser confocal culture dish, washed in PBS and fixed in 4% methanol for 30 min. After DAPI staining for 5 min and PBS washing for 5 min × 4. Fluorescent signals were visualized using an Olympus confocal laser scanning microscope (FV3000). TD channel represented the white light.

### Measurement of emission and excitation spectrum

GBM tissues were cut into small pieces and placed in a 2 ml EP tube, which were homogenated in 1.5 ml of RIPA lysate with 1.5 mM EDTA and placed in the 4ºC refrigerator in the dark for 30 min. After centrifugation at 12,000 rpm for 5 min, the supernatant was stored at -20ºC in the dark. The mixture containing 50 µl of the protein supernatant and 50 µl of PBS was subjected to the measurement of emission and excitation spectrum by the microplate reader (Synergy H1, BioTek).

### Flow cytometry

Cells were digested in trypsin and collected in a 1.5 ml EP tube. U87 and U87-GFP cells were considered as negative and positive control respectively. Optical density at 405 nm and 488 nm wavelength was measured by flow cytometry.

### FRET assay

HFP1 and HFP2 were labeled as the acceptor and donor fluorophore, respectively. Four randomly selected fields per sample were observed for detectable light emission signals. The occurrence of FRET was determined after photobleaching, and the binding constant was calculated.

### Measurement of chemical properties of HFPs

50 µl of protein samples were subjected to water bath at a certain temperature for 20 min, followed by mixing with 50 µl of PBS in the dark. The pH level gradient was prepared by the mixture of HCl and NaOH, and the total volume was identical by adding ddH2O. In addition, 50 µl of protein samples were mixed with 50 µl of SDS solution, and those mixed in 50 µl of PBS were considered as controls. The emission and excitation spectrum were measured by the microplate reader (Synergy H1, BioTek).

### Statistical analyses

Data were expressed as mean±standard deviation from three replicates. Differences between groups were compared by the Student’s t test. Statistical analyses were performed by SPSS 21.0, and P<0.05 considered as statistically significant.

## Results

### HFP1 is identified in primary GBM cells

primary GBM cells from clinical specimens of 6 GBM patients were isolated and cultured (Fig. SA). Confocal microscopic images of primary GBM cells were captured at 488 nm, 555 nm and 633 nm wavelength, respectively (Fig. 1A and B). Cross-over or bleed-through of fluorescence emission was prevented. Interestingly, an unknown fluorescent protein excited by 488 nm wavelength was detected, whilst no emissions were detected at the same intensity in the other two channels (Fig. 1A and B). We therefore nominated the unknown fluorescent protein as the human fluorescent protein Ⅰ (HFP1), which required a higher laser intensity than that of DAPI labeling. HFP1 was stained in a spotty shape (Fig. 1A).

**Figure 1.**
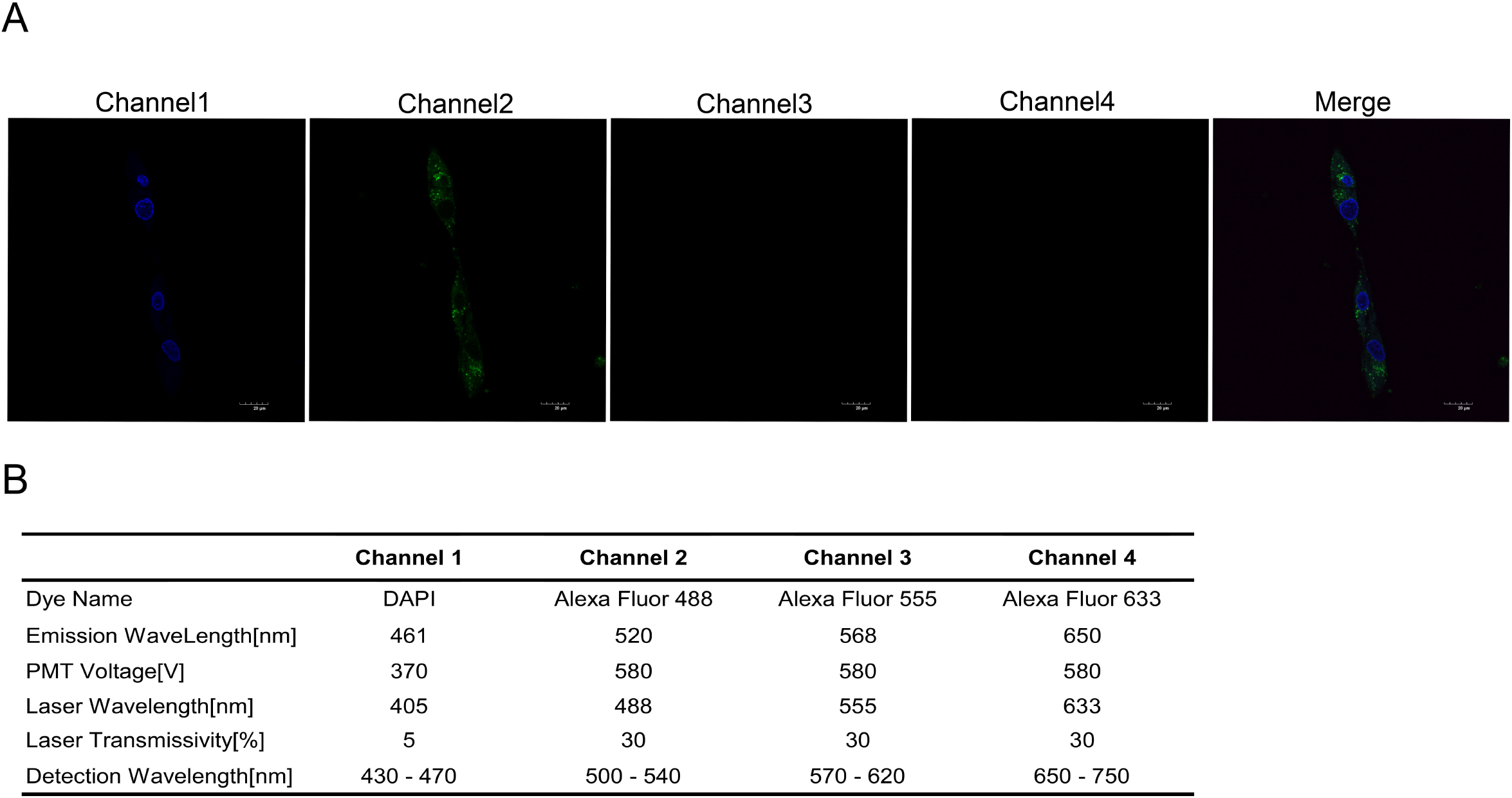
A. Confocal microscopic image of primary GBM cells. Cell nuclei were stained by DAPI. Scale bar = 20 µm. B. Scanning parameters of confocal microscopy.

### HFP2 is expressed in primary GBM cells and forms a polymer with HFP1

To exclude the possibility that the fluorescent signal of HFP1 was induced by the DAPI labeling, cell nuclei were not stained by DAPI and the cells were localized with light microscopy (TD channel). All fluorescent signals in the cytoplasm of primary GBM cells were detected at the 405 nm wavelength (Fig. 2A and B). It is shown that there was an unknown fluorescent protein that could be excited by 405 nm, which was nominated as HFP2 (Fig. 2A and B). A higher resolution ratio was adopted for enhancing the accuracy of fluorescence staining. As shown in the merged image, HFP1 and HFP2 were highly co-localized in primary GBM cells. Inspired by a previous study about the Aequorin and wtGFP, we speculated that HFP1 and HFP2 exerted a synergistic effect by forming a polymer, because they were both mainly distributed in the cytoplasm, rather than the nuclei.

**Figure 2.**
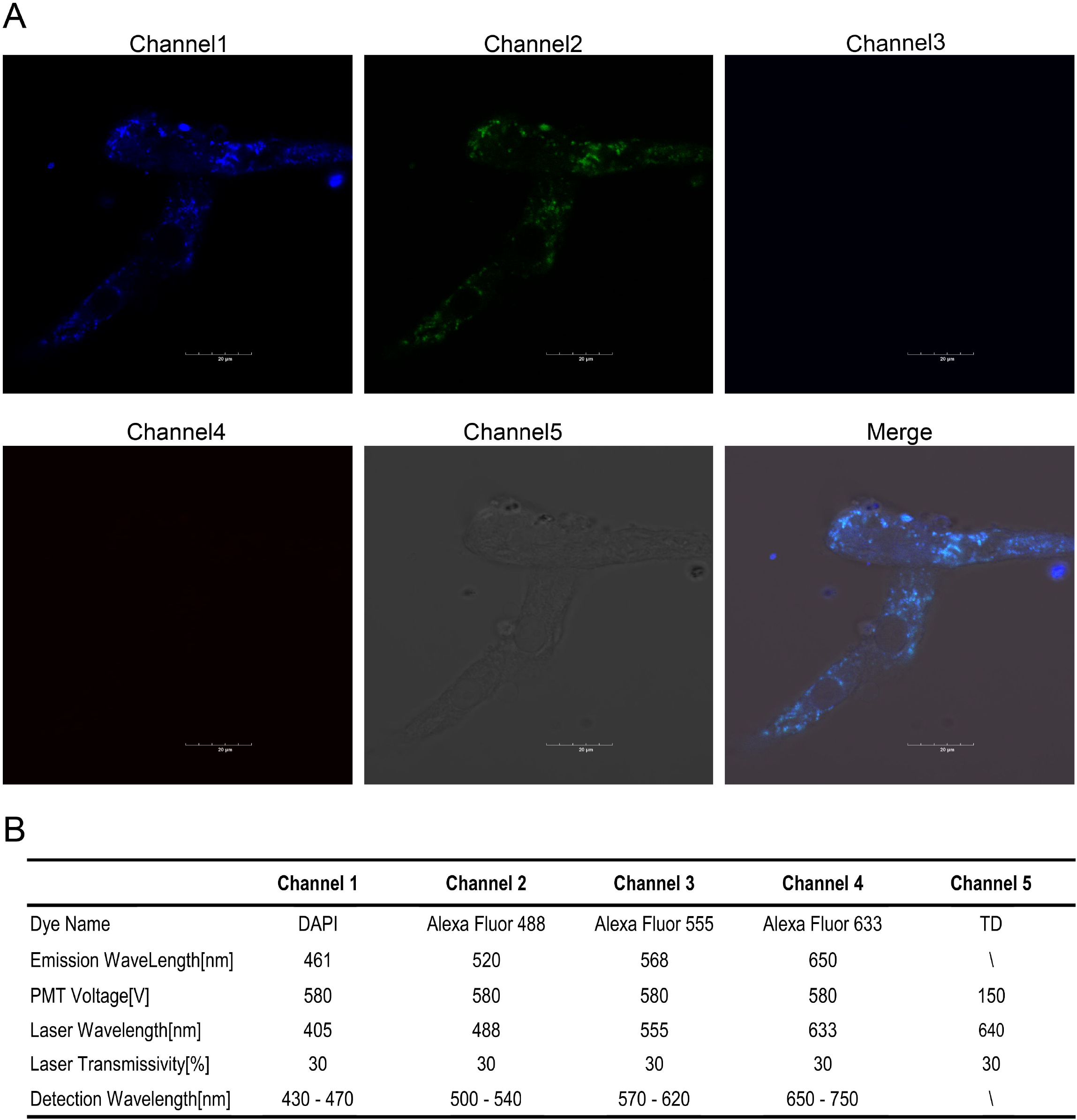
A. Confocal microscopic image of primary GBM cells. Cell nuclei were not stained. Scale bar = 20 µm. B. Scanning parameters of confocal microscope.

### The maximum emission and excitation spectrum of HFP1 and HFP2, and the identification of the FRET

Through measuring the emission and excitation spectrum of HFP1 and HFP2 collected from 20 GBM samples, it is found that the maximum emission and excitation wavelength of HFP1 was 542 nm and 467 nm, respectively, which was 448 nm and 378 nm in HFP2 (Fig. 3A and B). In addition to an emission peak at 448 nm wavelength, another emission peak at 540 nm was discovered in HFP2 when the spectrum was expanded to 600 nm wavelength. Interestingly, the maximum emission peak of HFP2 at 540 nm was spectrally overlapped with the maximum emission wavelength of HFP1, suggesting the presence of FRET (Fig. 3A and B). However, we did not detect relevant fluorescent signals in protein samples collected from U87, U251, 98T and GES-1 cell line (Fig. 3C). To further identify the distance-dependence interaction between HFP1 and HFP2, FRET assay was performed to directly detect the oligomerization state. After photobleaching in HFP1, the fluorescence intensity of HFP1 significantly decreased, while that of HFP2 slightly increased, suggesting the partial energy transfer from HFP2 to HFP1 (Fig. 3D). The binding constant of HFP1 and HFP2 was 20.3%.

**Figure 3.**
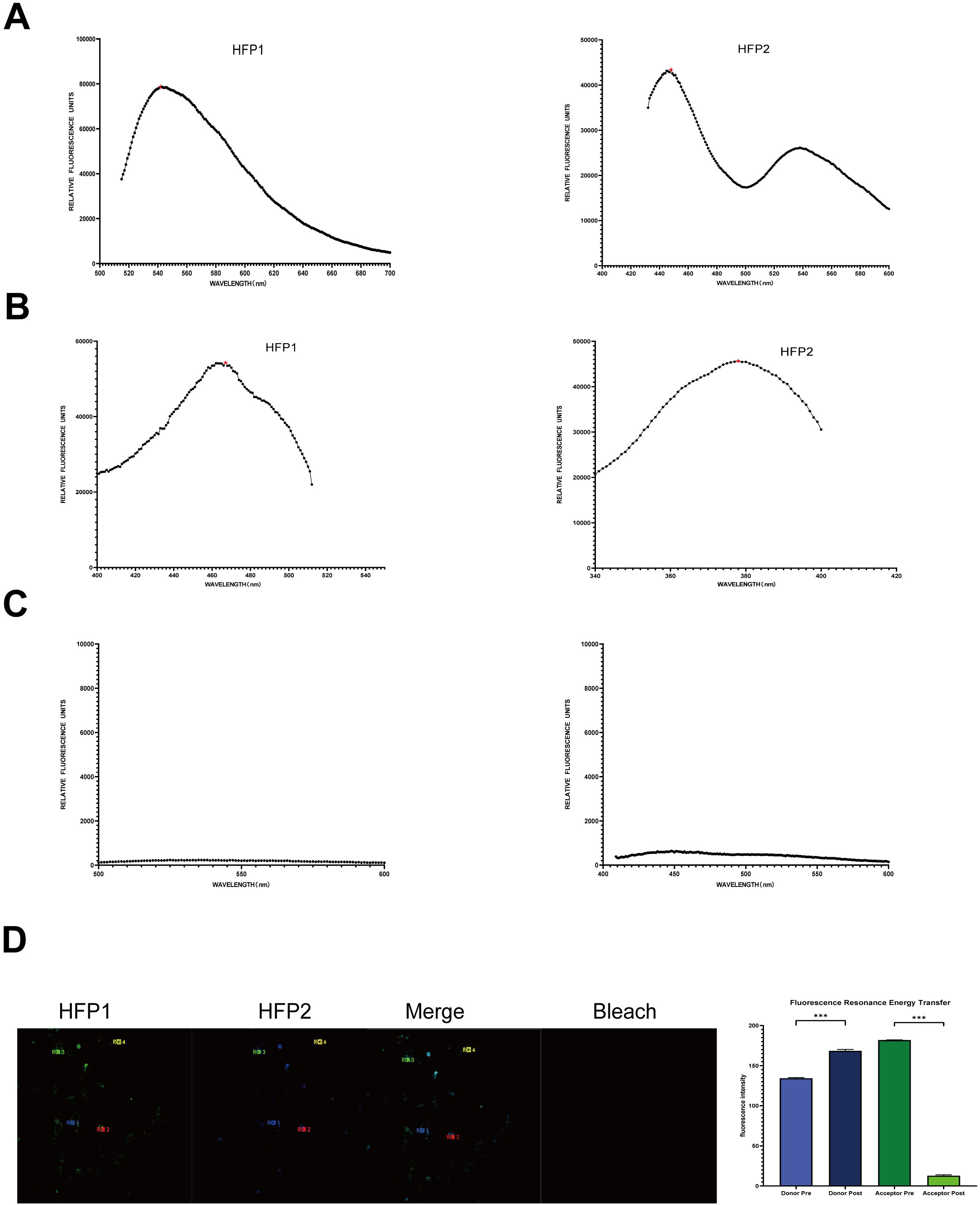
A. The emission spectrum of HFPs. B. The excitation spectrum of HFPs. C. The emission spectrum of HFPs in GBM cell lines. D. FRET assay of primary GBM cells before and after photobleaching. Data are presented as the mean ± SEM of three independent experiments. Significant results are presented as \**P<*0.05, \*\**P<*0.01, and \*\*\**P<*0.001.

### Identification of HFP1 and HFP2 in human GBM specimens

HFP1 and HFP2 were detected in human GBM specimens collected from the surgery, primary GBM cells, U87 cells (negative control) and GFP stable cell line-U87 (U87-GFP, positive control) by flow cytometry. HFP1 and HFP2 signals were both detected in tissue and cell samples of GBM, suggesting the positive expressions of HPFs in GBM (Fig. 4A& Fig. SB). Notably, we also detected HFP1 and HFP2 in 3 clinical samples of human cerebral trauma, and only HFP2 was detectable (Fig. 4A& Fig. SB). It is indicated that there may have normal cells expressing HFP2 in the human brain. Furthermore, the chemical properties of HFP1 and HFP2 were assessed. At the temperature of 60ºC, the activity of HFP1 reached the peak (Fig. 4B). HFP1 activity peaked in the environment of pH 6, which gradually decreased. The activity of HFP2 increased with the pH level and peaked at pH 12, which suddenly decreased in the environment of pH 13 (Fig. 4C). In addition, the influence of sodium dodecyl sulfate (SDS) solution on HFP activities was explored, which resulted in the loss of fluorescence activities of both HFP1 and HFP2 by denaturation (Fig. 4D).

**Figure 4.**
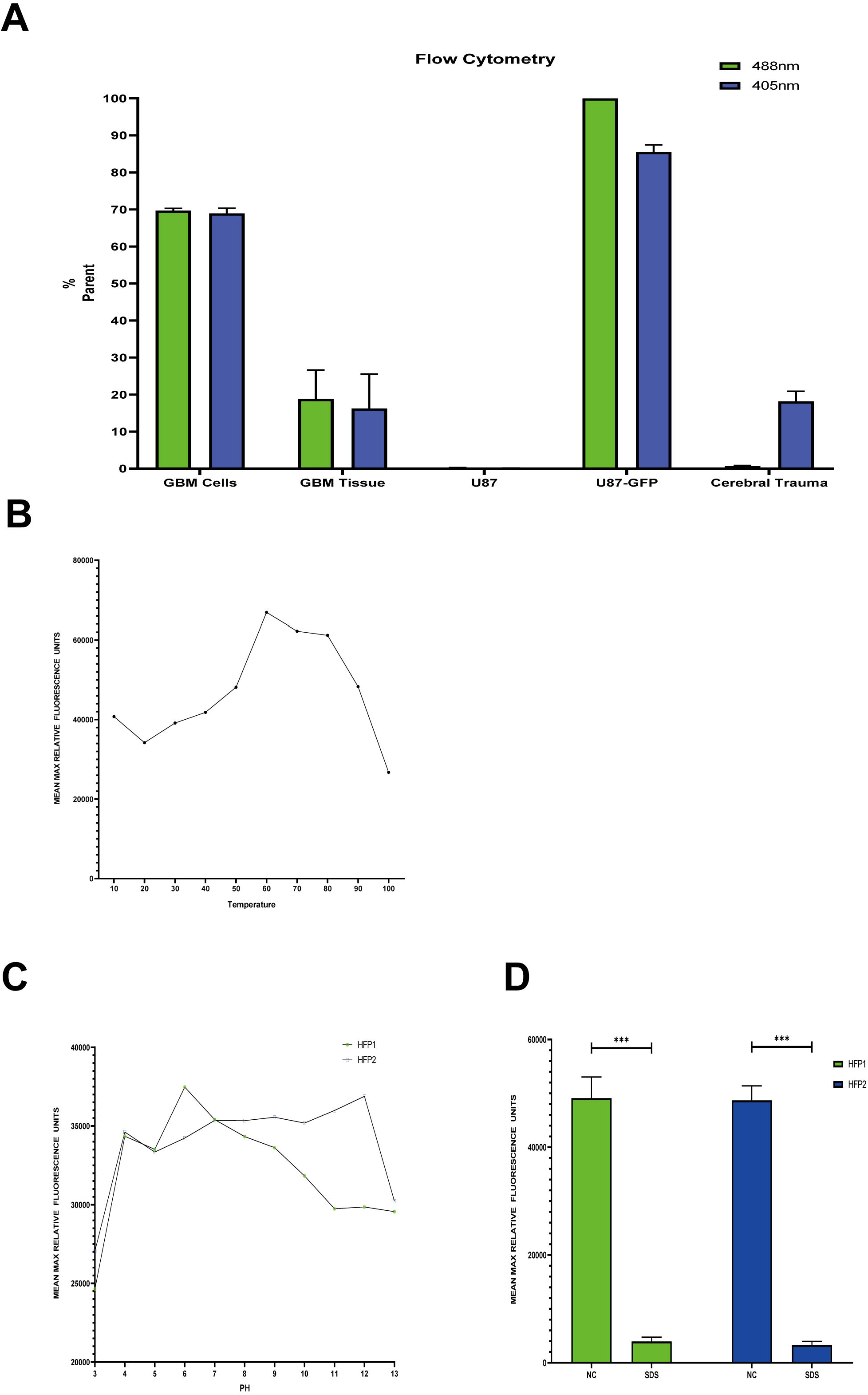
A. Flow cytometry analysis of GBM tissues, primary GBM cells, U87 cells, U87-GFP cells and cerebral trauma samples. B. The emission and excitation spectrum of HFPs responsive to the temperature change. C. The emission and excitation spectrum of HFPs responsive to the pH level change. D. The emission and excitation spectrum of HFPs responsive to SDS solution. Data are presented as the mean ± SEM of three independent experiments. Significant results are presented as *P<0.05, **P<0.01, and ***P<0.001.

## Discussion

This study for the first time discovered 2 novel fluorescent proteins HFP1 and HFP2 in primary GBM cells, which was of great significance in the medical research. GBM is deeply located in the brain, and the fluorescent signals of GBM cells are difficult to be excited by the conventional fluorescence channels. Based on previous immunofluorescence staining experiences, HFP1 was more difficult to be excited than commonly used fluorescent conjugate secondary antibodies, and its fluorescence intensity was relatively weak. The polymer formed by HFPs may be essential for polymerization energy.

In the present study, we identified the genetic information of fluorescent proteins in the human genome, and highlighted the potential functions of HFPs in GBM cells. Unfortunately, DNA sequences, more physical and chemical properties and crystal structures of HFP1 and HFP2 failed to be determined, which will be explored in the future. Moreover, positive and negative influences of ions on HFPs were unknown. Due to the long-term cell culture and cell passage, expression levels of HFPs in in vitro cultured cell lines are far lower than those in primary cells, although the isolation and culture of primary GBM cells are challenging and expensive, and their use is also restricted by the limited human samples.

The unknown type of a protein itself is controversial. Based on the current findings, the newly discovered proteins were determined as HFPs, rather than other types of organic compounds. Existing evidences have shown that similar genetic sequences to those of GFP-like proteins in human genome are not existed^1^. However, our data of clinical samples of human cerebral trauma revealed that cells only expressing HFP2 may exist in human brain tissues. The conventional methods are unable to determine the structure and functions of HFPs, which can only be solved by quantum mechanics^1^.

The desire to survival is the top priority of living things. Environmental factors influence all types of organisms, and the evolution is the responsiveness of genetic information to environmental changes. So far, the macroscopic and the microscopic world cannot be unified by a consensus evolutionary theory. Here, we speculated a universal evolutionary theory that was applied to various environmental changes, in which the macroscopic world was the consequence of repeated evolutions of the microscopic world to be strictly performed. According to the assumed theory, the inorganic telescope in the macroscopic world is a repeated evolution from eyes, and in parallel, the inorganic microscope may also be the repetition of some organic functions of cell bodies. The light source is essential for the microscopy, and thus, there may have potential sensitive light sources in cells. Previous results have shown that GBM cells share a similar human genome to that of normal somatic cells. Therefore, we searched potential build-in light by a direct excitation in primary GBM cells. At present, which types of somatic cells positively expressing HFPs are unknown. Considering that fluorescent proteins are the only fluorescent probes from the natural source, their potential biological functions in human bodies are of great significance to be further explored.

Proteins that interact with HFPs are very likely to be existed, and the corresponding signals can be identified. According to the law of minimum energy in living things, light signals produced by the human eyes and the microscope in cells may be similarly processed. If the crystal structure, and the physical and chemical properties of HFPs can be successfully identified, they may serve as a novel non-invasive medical detection method. The in vivo optical detection will astonishingly advance the progression of human medicine. The unified evolutionary law that living things ultimately conform is worthy of further exploration. Searching for a mathematical form to prove that genetic information itself may also be a certain form of particles is of great significance. Owing to the basic properties of life particles, they can only be distributed in DNAs or RNAs, rather than proteins. We believed that electromagnetic signals in the environment may directly affect genetic information, which is reflected as the adaptation of living things to the change of the environment.

## Supporting information

support table

support figure

## Acknowledgement

Not applicable.

