## Supplementary material for "The identification of two newly discovered fluorescent proteins in human glioblastoma": support table

**Table S1: Summary of clinical GBM patients**

| Characteristic | All patient High CircASAP1 expression (n=26) |
|---|---|
| **Type** | |
| Pri GBM | 15 |
| Rec GBM | 11 |
| **Sex (n)** | |
| Male | 16 |
| Female | 10 |
| **age** | |
| ⩾45 | 19 |
| ＜45 | 7 |
| **Tumor location** | |
| Frontal | 11 |
| Non-frontal | 15 |
| **KPS score** | |
| ≥80 | 15 |
| ＜80 | 11 |
| **MGMT promotor status** | |
| Methylated | 18 |
| Unmethylated | 8 |
| **Extent of surgery** | |
| Total | 19 |
| Subtotal | 7 |
| **IDH1/2 genotype** | |
| Mutation | 5 |
| Wild-type | 21 |

**Abbreviations**: KPS, Karnofsky performance status; MGMT, O-6-methylguanine-DNA-methyltransferase; IDH1/2, isocitrate dehydrogenase 1 and 2.
