## Supplementary figures and images for "The identification of two newly discovered fluorescent proteins in human glioblastoma"

### support figure

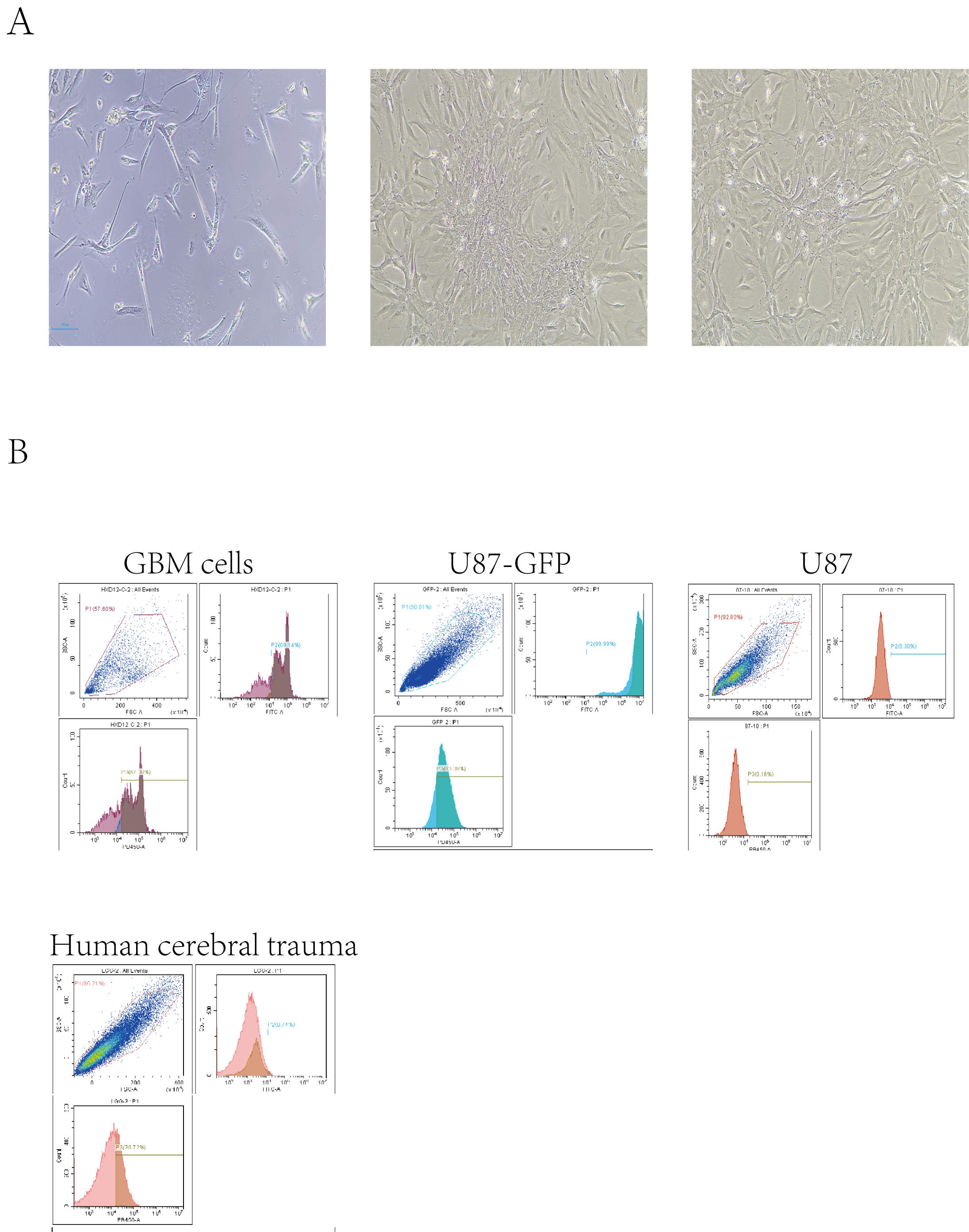
